# Attention Towards Emotions is Modulated by Familiarity with the Expressor. A Comparison Between Bonobos and Humans

**DOI:** 10.1101/2020.05.11.089813

**Authors:** Evy van Berlo, Thomas Bionda, Mariska E. Kret

**Affiliations:** Leiden University, Institute of Psychology, Cognitive Psychology Unit, Leiden, The Netherlands; Leiden Institute for Brain and Cognition (LIBC), Leiden, The Netherlands; Apenheul Primate Park Foundation

**Keywords:** AFFECT, GREAT APE, INTERGROUP BIAS, AFFECTIVE ATTENTION, SOCIAL PROCESSING

## Abstract

Why can humans be intolerant of, yet also be empathetic towards strangers? This cardinal question has rarely been studied in our closest living relatives, bonobos. Yet, their striking xenophilic tendencies make them an interesting model for reconstructing the socio-emotional capacities of the last common ancestor of hominids. Within a series of dot-probe experiments, we compared bonobos’ and humans’ attention towards scenes depicting familiar (kith and kin) or unfamiliar individuals with emotional or neutral expressions. Results show that attention of bonobos is immediately captured by emotional scenes depicting unfamiliar bonobos, but not by emotional groupmates (Experiment 1) or expressions of humans, irrespective of familiarity (Experiment 2). Using a large community sample, Experiment 3 shows that human attention is mostly captured by emotional rather than neutral expressions of family and friends. On the one hand, our results show that an attentional bias towards emotions is a shared phenomenon between humans and bonobos, but on the other, that both species have their own unique evolutionarily informed bias. These findings support previously proposed adaptive explanations for xenophilia in bonobos which potentially biases them towards emotional expressions of unfamiliar conspecifics, and parochialism in humans, which makes them sensitive to the emotional expressions of close others.

## Introduction

Emotional expressions are a major force in navigating the social world; they provide valuable insights into the emotional states of others and help to predict others’ future behaviors^1^. Bonobos (*Pan paniscus*) have been put forward as a model for studying the phylogeny of human emotion processing ^2–4^ because of their striking xenophilic tendencies ^3,5–7^. They also form an interesting comparison species for gaining evolutionary insights, on the one hand, into humans’ empathy towards strangers ^8^, and parochialism on the other ^9,10^. However, we currently have only limited knowledge about how bonobos process emotional expressions. Here, we investigate how bonobos compared to humans, process the emotional expressions of familiar and unfamiliar others.

Over evolutionary time, selective pressures gave rise to brains that are able to quickly attend to and understand emotional expressions in order to facilitate communication between individuals ^11^. Research in humans has demonstrated that already during the earliest stages of visual perception, attention is attuned to emotional expressions ^12,13^. Specifically, both threatening and positive signals in the environment can rapidly capture attention ^14^, and this attentional attunement is driven by both arousal-eliciting characteristics of the signal as well as its significance to the observer ^15,16^. Interestingly, a similar capacity has been observed in bonobos, humans’ closest living relatives ^2^. In an experimental setting, bonobos showed an attentional bias for emotional scenes depicting unfamiliar conspecifics, especially when these scenes were emotionally intense. This finding suggests that the attentional mechanisms that guide social perception have an evolutionarily old foundation, and were likely already present in the last common ancestor of *Pan* and *Homo*.

Emotions are most frequently expressed in the proximity of group mates, such as kin and friends. This is beneficial, because individuals rely on each other to attain personal goals in the short and long term ^17^. As attention gates which signals from the environment are preferentially processed, it is plausible that evolution fine-tuned this mechanism to quickly differentiate not only between emotional and neutral cues, but also between expressions of familiar, socially close group members and unfamiliar others. Compared to other great apes and humans, bonobos are strikingly xenophilic. Intergroup encounters in the wild proceed relatively peacefully, and neighboring groups have been observed foraging together ^18^. Furthermore, in experimental settings they show a prosocial preference for unfamiliar individuals rather than group members ^19^. In stark contrast, humans tend to prioritize their own group members over unfamiliar individuals when it comes to sharing resources^10^.

It has been argued that bonobos evolved into a more tolerant ape due to a relatively stable environment that reduced feeding competition. As a result, bonobos are able to form stable social parties in which females form alliances and reduce male aggression. Moreover, these stable social groups prevent extreme territorial encounters with other groups, leading to intergroup tolerance ^7,20^. Interestingly, the evolutionary environment of bonobos contrasts with that of humans. For substantial periods of time, ancestral humans migrated great distances across the globe as a result of the extraordinarily volatile climate that caused scarcities in resources. This paved the way for intergroup conflicts among our hunter-gatherer ancestors ^21^. In turn, these aggressive interactions have fostered a strong focus on the in-group (e.g. family and friends) on the one hand, and xenophobia on the other ^9^. At the heart of interactions with both strangers and socially-close others lie emotional expressions, as they communicate emotional states and intentions. As such, the discrepancy between how bonobos and humans evolved to interact with others offers a highly interesting motive for taking a closer look at how the two species process emotions of family, friends, and strangers. Specifically, we ask how familiarity impacts early attentional mechanisms that help distinguish between emotionally relevant signals from group members or other, unfamiliar individuals.

To make inter-species comparisons of selective attention for emotions possible, the emotional dot-probe paradigm has been proven useful ^22,23^. In the task, individuals have to press a central dot, followed by a short presentation of an emotional and a neutral stimulus. Another dot (i.e. the probe) then replaces either the emotional or neutral stimulus. Individuals are generally faster at tapping the probe that replaces the stimulus that immediately caught their attention (usually the emotional stimulus) compared to a probe replacing the other stimulus (e.g. the neutral stimulus). As such, the emotional dot-probe task provides an easy way to tap into the underlying attentional mechanisms that guide emotion perception.

In the current study, we investigate how bonobos and humans attend to expressions of emotion of familiar and unfamiliar individuals. Here, we define familiarity by the social and familial relationship between the observer and the expressor of emotions on the one hand, and unfamiliar others on the other. First, we investigate whether bonobos have an attentional bias towards emotional expressions of unfamiliar and familiar conspecifics (Experiment 1), followed by whether this bias extends to unfamiliar and familiar human expressions (Experiment 2). In Experiment 3, using a large community sample of zoo visitors, we investigate whether attention is attuned to emotional expressions of familiar (family and friends also visiting the zoo) or unfamiliar (other zoo visitors) people.

We hypothesize that bonobos will show an attentional bias towards emotions expressed by unfamiliar conspecifics ^2^ and that this bias will be dampened when seeing familiar conspecifics. Furthermore, since certain aspects of emotion processing are shared between humans and extant apes ^24^, we further predict that bonobos will show an attentional bias towards emotional expressions of humans. Whether this bias in bonobos is modulated by the familiarity of the human expressor is an exploratory question. For humans, we hypothesize that an attentional bias towards emotions exists for expressions of unfamiliar individuals ^22^. We also expect that this bias will be more pronounced for familiar individuals as compared to unfamiliar individuals, reflecting the parochial tendencies of humans ^9^. Finally, since attention is typically captured by biologically relevant and salient signals from the environment, we expect that in all three experiments, the more emotionally intense an expression is, the faster it will capture attention ^2^.

## Method

### Experiment 1: Bonobos’ attentional bias towards emotions of conspecifics

#### (a) Participants

Four female bonobos living in a social group of 12 individuals at Apenheul primate park in Apeldoorn, The Netherlands, took part in the study and were tested over a period of 4.5 months. Testing took place in the presence of non-participating group members and during winter when the park was closed for visitors. Testing took place three to four times a week in one of the indoor enclosures, and a session lasted roughly 15-20 minutes per individual.

Tests with the bonobos were conducted adhering to the guidelines of the EAZA Ex situ Program (EEP), formulated by the European Association of Zoos and Aquaria (EAZA). Bonobos participated voluntarily and were never separated from their group during testing. Only positive reinforcement (apple cubes) was used during training and testing, and each bonobo (including ones that were not tested) received an equivalent reward. Non-participating bonobos were distracted by the animal caretaker who conducted a body-part training task used for veterinary purposes (see Supplements for additional information).

#### (b) Apparatus

The experiment was conducted using *Presentation* (NeuroBehavioralSystems) on an Iiyama T1931SR-B1 touchscreen (19”, 1280×1024 pixels, ISO 5ms) encased in a custom-made setup (Figure 1). To limit exposure to the experimenter, rewards for correct responses were automatically distributed using a custom-made auto-feeder apparatus that dropped apple cubes into a funnel that ended underneath the touchscreen for the bonobo to grab. This allowed us to limit the amount of disruptions during testing. Bonobos were filmed from the side during all test sessions. These videos were later used to code behaviors during the task and filter out unsuccessful trials.

**Figure 1.**
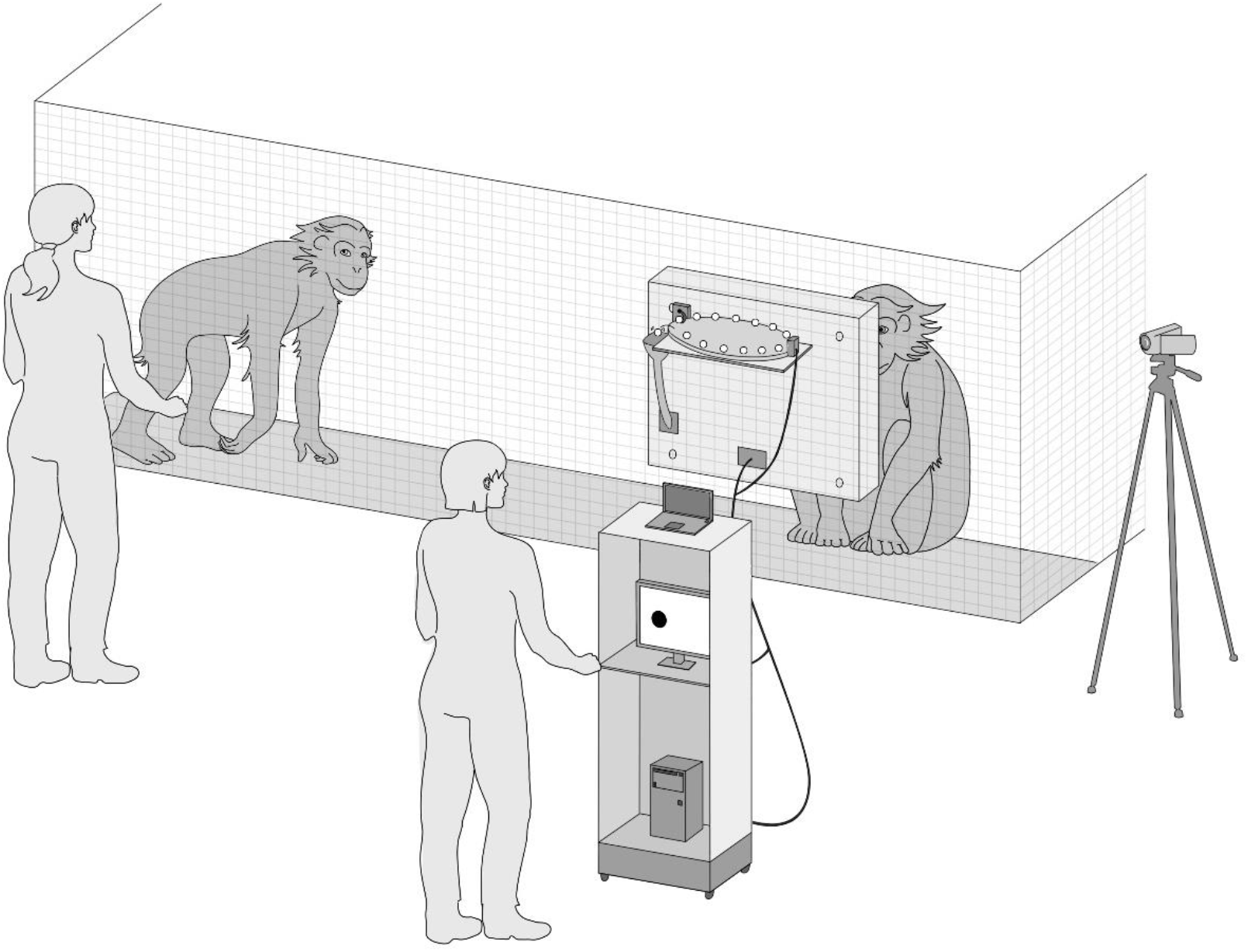
Abstract representation of the bonobo setup. The experimenter (right) controlled the experiment from behind the bonobo setup while a caretaker (left) distracted the other bonobos.

#### (c) Stimuli and validation

Stimuli consisted of bonobo pictures collected in different zoos and from the internet. Stimuli of familiar individuals consisted of pictures of the group living in Apenheul. In total, the study included 656 unique pictures (346 of familiar and 310 pictures of unfamiliar individuals) (see Supplements and Table S1). All pictures were resized to 330 x 400 pixels and showed either a neutral scene (i.e. individuals sitting or lying down or involved in a non-social activity, showing a neutral expression) or an emotional scene. Emotional scenes depicted individuals in distress, playing, grooming, having sex, yawning, or scratching (Table S2 and Figure S1). Scratching is indicative of stress in both primates and humans ^25^ and by using it, we increased the number of negatively valenced stimuli. While yawning is not an emotion in itself, it is a highly contagious behavior that can be viewed as a proxy for empathy (but see ^26^) and it captured attention faster than other categories in the study by Kret et al. (2016). We matched emotional and neutral scenes on the number of individuals depicted, their identity, and by visual inspection of color and luminance. All pictures were rated on emotional valence and intensity by three primate experts from Apenheul and three primate researchers, who showed high intraclass correlations (ICC_valence_ = .82, ICC_intensity_ = .87, (see Supplements and Table S3).

#### (d) Procedure

The bonobos were already familiarized with the dot-probe procedure during a previous study ^2^, but did go through a training phase (see Supplements). In the experiment, a trial started with the presentation of the start dot in the middle, lower part of the screen (Figure 2). After the bonobo pressed the dot, a neutral and an emotional stimulus appeared on the left and right side of the screen for 300ms. Stimuli were always either of bonobos familiar to the participant or of unfamiliar individuals (thus, we never combined e.g. an emotional picture of a familiar with a neutral picture of an unfamiliar or vice versa). Stimuli were subsequently followed by another dot (the probe) replacing either the neutral or emotional stimulus. The probe remained on the screen until touched, after which an apple cube was provided through the auto-feeder system. After a delay of 2000ms the next trial started.

**Figure 2.**
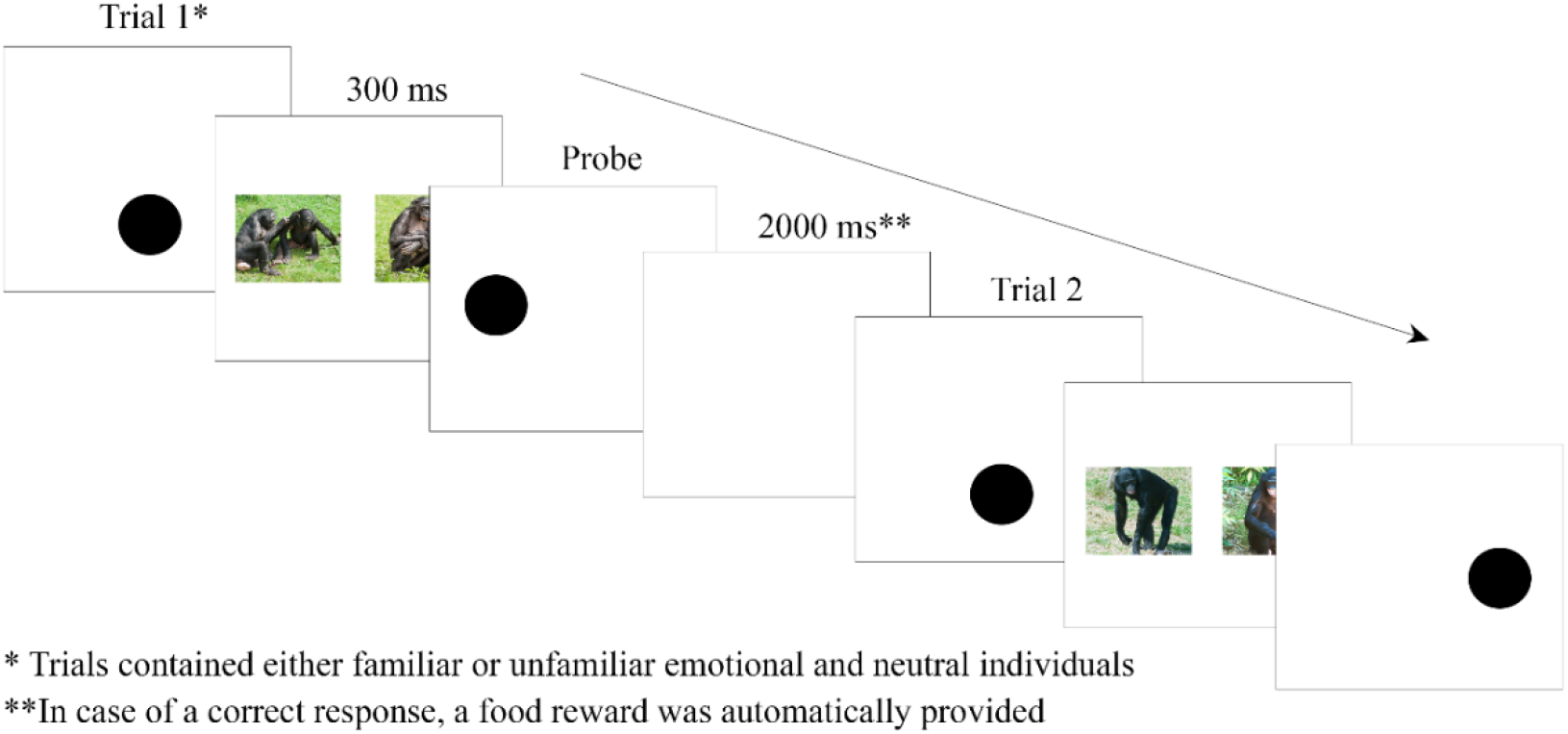
Trial outline of the bonobo dot-probe task.

Each test session consisted of 25 trials in which the location of the stimuli on the screen (i.e. left/right) and the location of the probe (i.e. behind the emotional or neutral stimulus) were counterbalanced, and the order of stimulus presentation was randomized based on emotion category and familiarity. In each session, half of the trials consisted of emotional and neutral stimuli of familiar individuals, and half of emotional and neutral stimuli of unfamiliar individuals. If a trial was deemed unsuccessful directly after testing, it was repeated. In total, each bonobo finished between 21 to 24 sessions and on average a total of 541 trials (*SD* = 28.76, Table S4 in Supplements).

#### (e) Data Filtering

Behaviors were scored from the videos by two experts with high agreement (ICC = .95, *p* < .001). Next, extreme reaction times (RT; 250 ms < RT < 5000 ms) were filtered out and trials where bonobos were not attending the task. Finally, trials with RTs higher than the median RT per subject minus 2.5 * the median absolute deviation per subject (MAD) were excluded. Based on this filter, 514 trials (23.8%) were removed from the analysis (see Supplements and Table S4).

#### (f) Statistical Analysis

We used generalized linear mixed models (SPSS v20, α =.05) for the analyses, which contained a nested structure defined by trials (25) nested within sessions (21-24) nested within participants (4). We used random intercepts per participant and participant*session. Reaction time was used as the dependent variable, and as they are typically skewed, we selected a gamma-distribution with a log-link function.

In the statistical model we included *Congruency* (i.e. the probe replaces an emotional [congruent] or neutral [incongruent] stimulus) and *Familiarity* (i.e. familiar versus unfamiliar bonobos), and their interaction terms as fixed factors. Next, to replicate findings by Kret et al. (2016) who showed that the more emotionally intense a stimulus is, the faster it captures attention, we calculated a difference score between the average emotional intensity for stimuli that were replaced by the probe (probe image) and the distractor image (nonprobe image), as the two images are competing for attention. Positive values meant that the probe image was emotionally more intense than the nonprobe image. In a second model, we then included *Intensity Difference Score* as a fixed factor to test its effect on reaction time.

### Experiment 2: Bonobos’ attentional bias towards emotions of humans

#### (a) Participants

See Experiment 1.

#### (b) Apparatus

See Experiment 1.

#### (c) Stimuli and validation

Stimuli consisted of isolated emotional and neutral human faces that were either unfamiliar to the bonobos (NimStim Set of Facial Expressions ^27^) or familiar (4 female bonobo caretakers that interact with the bonobos on a daily to weekly basis, and 2 female experimenters that recently trained and tested the bonobos). Emotional expressions consisted of six basic human expressions ^28^: anger, fear, happiness, sadness, surprise, and disgust. (Supplements Figure S2). Stimuli were in color and sized 330×400 pixels. In total we had 144 stimuli (72 of familiar and 72 of unfamiliar individuals).

To check the validity of our stimulus materials, we first asked an independent group of research assistants (N = 5) to rate the materials on emotion type (whether the stimulus is an emotional or neutral expression), arousal and authenticity. Results indicated the following intraclass correlations: intensity of the stimuli (ICC = .78), emotion (ICC = .66), and authenticity (ICC = .69; see Table S7a-b in Supplements). Furthermore, as a control measurement, we had a group of zoo visitors (N = 150, M_age_ = 39.97, SD = 14.98) take part in a dot probe experiment using the Caretaker and NimStim stimuli (see Supplements).

#### (d) Procedure

The procedure for bonobos in Experiment 2 was similar to Experiment 1, except that human stimuli were used. In total, each bonobo finished 345 trials on average (*SD* = 24.56), divided over 13-15 sessions per individual (Table S8).

#### (e) Data Filtering

As in Experiment 1, two experts rated the videos in high agreement (ICC = .96, *p* < .001). We used the same data filters as in Experiment 1, resulting in removal of 373 trials (27.1%, Table S8).

#### (f) Statistical Analysis

Similar to Experiment 1, we used a GLMM with a nested structure with trials (25) nested within sessions (13-15) nested within participants (4) and random intercepts per participant and per participant*session. The dependent variable was reaction time using a gamma-distribution with a log-link function. We included *Congruency*, *Familiarity* (familiar versus unfamiliar human model) and their interaction terms as fixed factors. As a control, a group of novel visitors who were not familiar with any of the individuals on the stimuli performed the same experiment, and using a GLMM, we tested for effects of *Congruency* and *Stimulus Set* (i.e. NimStim and Caretaker) and their interaction terms on reaction time.

### Experiment 3: Humans’ attentional bias towards emotions of conspecifics

#### (a) Participants

Participants in the dot-probe task (N = 449, 196 men) were adults and children visiting primate park Apenheul in Apeldoorn, The Netherlands. Individuals were between 3 and 84 years old (*M* = 24.9, *SD* = 16.43). Apenheul allowed us to set up a research corner close to the bonobo enclosure where we could test the visitors (Figure 4). We only approached individuals who were visiting the zoo with at least one other person. To keep the human and bonobo experiments as similar as possible, we selected individuals based on their relation to each other (the “familiar” condition of the bonobos only contains group members, either kin or friends, see Table S11a-b in Supplements).

#### (b) Apparatus

Participants performed the experiment on an Iiyama T1931SR-B1 touchscreen (19”, 1280×1024 pixels, ISO 5ms) using E-Prime 2.0. The tests were conducted in an indoor compound in which visitors of the zoo could see the bonobos. The touchscreen was placed on a table and participants were seated with their back against a wall to prevent others from distracting them (Figure 3).

**Figure 3.**
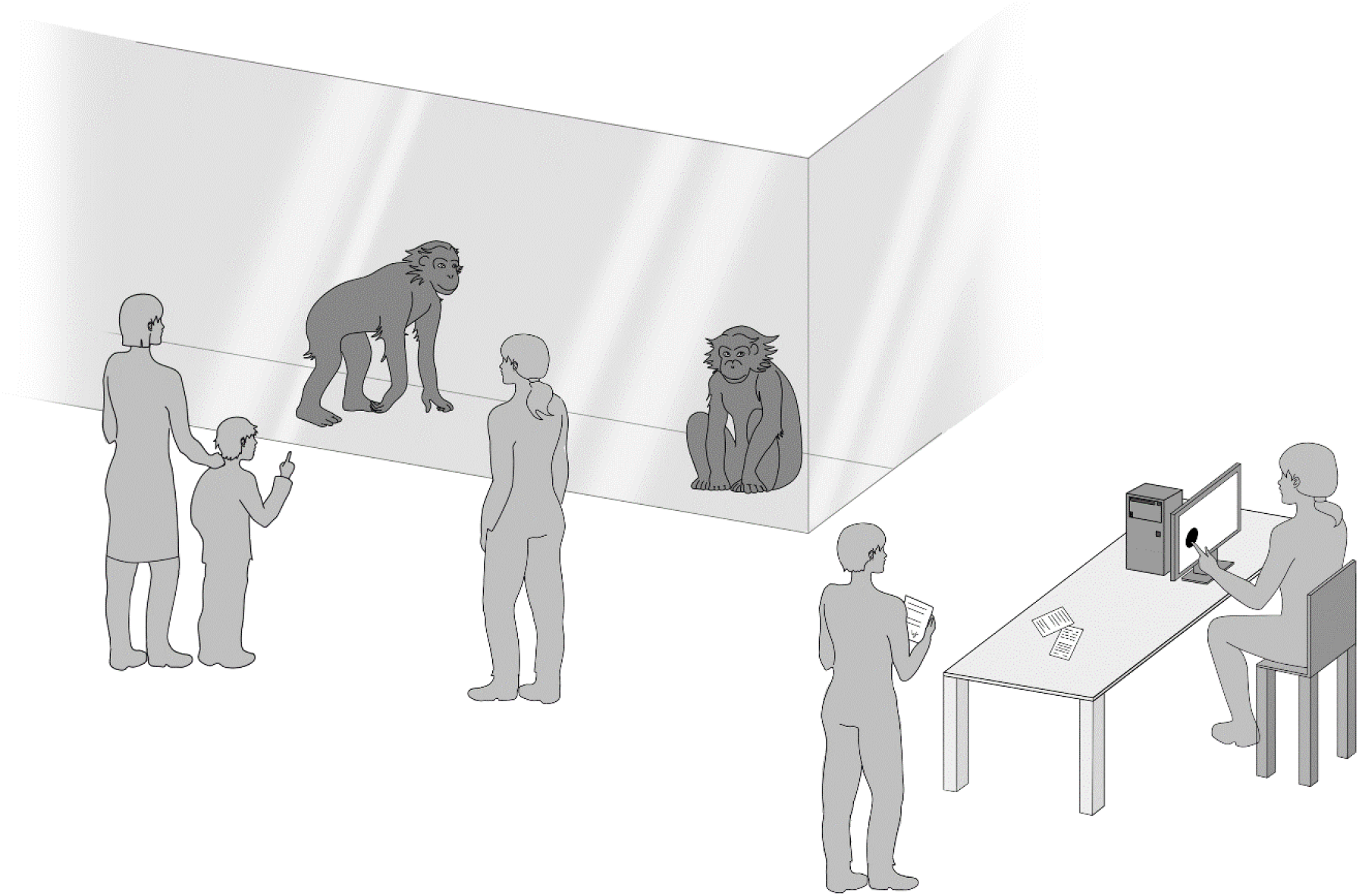
Abstract representation of the human setup near the bonobo enclosure.

#### (c) Stimuli and validation

Stimuli consisted of pictures of the face showing either an emotional (angry, fearful, happy, sad) or neutral expression presented against a neutral background. Each stimulus showed either a familiar individual (a family member or a close friend or colleague), or an unfamiliar individual (a previous participant in the study). We used four out of the six basic emotions ^28^ for practical reasons; the task would become undesirably long given that our participants were voluntarily taking part in our study. Pictures were sized 400×300 pixels. Each participant completed 40 trials (i.e. 20 trials showing emotional and neutral pictures of familiar individuals). The number of stimuli per emotional category was counter-balanced across participants, and stimulus combinations (emotional plus neutral) were presented in a semi-randomized order.

A total of 4040 pictures were split into three sets and rated on intensity, emotionality (whether a stimulus depicts an emotional or neutral expression), and authenticity by 18 university graduate and PhD candidates, and on average there was good agreement (ICC_intensity_ = .80, ICC_emotion_ = .80, and ICC_authenticity_ = .68, see Supplements: Table S12a-d).

#### (d) Procedure

Zoo visitors passing by the bonobo enclosure with at least one other person were approached by test leaders. If they wanted to participate, the experimenter decided which participant was going to perform the dot-probe task (i.e. dot-probe participant) and who was going to be on the photos that subsequently served as stimulus material (i.e. photo participant). Photo participants never performed the dot-probe task, and dot-probe participants were never photographed. Individuals could only participate in the study once.

After reading the information sheet and signing a consent form, photos were made of the photo participant. The procedure for photographing participants was similar to the procedure for the control experiment in Experiment 2 (but see Supplements).

Next, the pictures were loaded into the software and the dot-probe participant was then seated behind the touchscreen, followed by the experimenter entering personal data (age, handedness, sex of both the dot-probe and photo participant, the nature of their relationship, and how often they see each other (Table S11a-b). The task was the same as for the bonobos, except no fruit rewards were provided (Figure S3). The task consisted of 40 trials that were presented in random order (i.e. 20 trials using stimuli of familiar individuals, and 20 of unfamiliar individuals). Furthermore, in both familiar and unfamiliar trials, participants saw five different stimuli of each of the four basic emotions (anger, fear, happiness, sadness). The location of the stimuli on the screen (left/right) and the location of the probe were counterbalanced, and stimuli were presented in a randomized order. The whole procedure took about 15 to 20 minutes to complete.

#### (e) Data filtering

We filtered reaction times (RTs) with extreme values (i.e. 250 ms < RT < 5000 ms). As our dataset contains a large age range, we also filtered reaction times per age category (0-5, 6-10, 11-15, …, 56-60, 61-85) and calculated the median absolute deviation for reaction times per age categories. Finally, we used the following filter: [RT < (Median RT + (2.5 * Median Absolute Deviation))]. In total, we excluded 14.63% of the data for further analysis.

#### (f) Statistical analysis

Data were analyzed using GLMMs in SPSS (v20, α = .05). In all models, experimental trials (40) were nested within participants (449). We used reaction time (ms) as the dependent variable and a gamma-distribution with a log-link function, and random intercepts for all participants. The first model included *Congruency*, *Familiarity*, and their interaction terms as fixed factors. Furthermore, as we expected that a bias for emotions is stronger when the expression is more intense, we calculated a difference score similar to what we calculated in Experiment 1 for bonobos. We then used *Intensity Difference Score* as a fixed factor to predict reaction time.

## Results

### Experiment 1: Bonobos’ attentional bias towards emotions of conspecifics

We aimed to replicate and extend previous findings by Kret et al. (2016) and tested for a possible interaction between familiarity and emotional attention in bonobos, and effects of intensity of stimuli on reaction time. In our first model, we found a significant interaction effect (*F*(1, 1644) = 4.65, *p* = .031); bonobos responded faster on probes replacing emotional (*M* = 516.40, *SE* = 58.40) rather than neutral scenes (*M* = 526.65, *SE* = 59.57) in the *Unfamiliar* condition ((*t*(1644) = 1.95, *p* = .052) but not in the *Familiar* condition, (t(1644 = −1.07, *p* = .285, Figure 4 and Table S5)^29^. In short, the attention of bonobos is immediately drawn to emotional scenes instead of neutral scenes of unfamiliar conspecifics, but not of familiar conspecifics.

**Figure 4.**
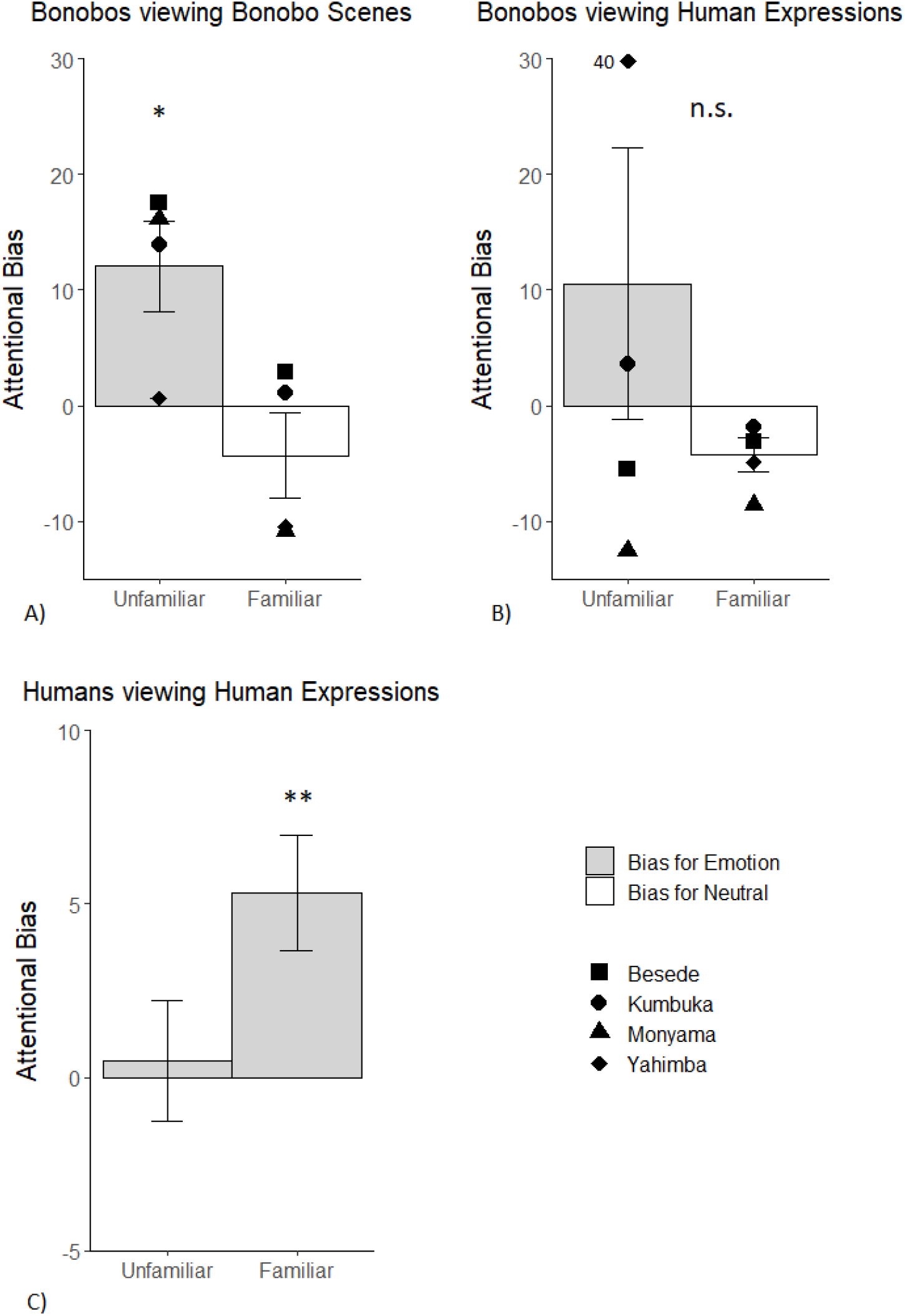
A) Bonobos show an attentional bias for emotions of unfamiliar, but not familiar conspecifics. B) Bonobos do not show an attentional bias for emotions of familiar and unfamiliar humans. The diamond-shaped symbol shows an individual score that falls outside of the graph range. C) Humans have an attentional bias for emotional expressions of familiar others. For visualization purposes, attentional bias was calculated as a difference score between mean reaction times (RTs) on emotional scenes minus mean RTs on neutral scenes per condition (Unfamiliar, Familiar). Bars in the positive direction indicate a bias towards emotional scenes (bonobos) or expressions (humans) rather than neutral scenes/expressions. Error bars represent the SEM. \**p* < .05.

In order to gain more insight into the attentional bias towards emotional expressions of unfamiliar individuals, we ran a second model with *Intensity Difference Score* as a fixed factor to specifically investigate whether emotional intensity of the stimulus would be positively associated with the observed attentional bias. As expected, reaction times were faster the higher the emotional intensity of the probe picture in relation to the non-probe picture (*F*(1, 859) = 8.38, *p* = .004, Table S6)^29^. As such, the more emotionally intense a signal of a conspecific is, the faster it captures attention in bonobos.

### Experiment 2: Bonobos’ attentional bias towards emotions of humans

To investigate whether bonobos’ attention for the emotional expressions of unfamiliar bonobos extends to human expressions, bonobos performed at dot-probe with familiar (i.e. zoo keepers) and unfamiliar (i.e. NimStim faces^27^) human emotional and neutral stimuli. Though the effect appeared to be in the expected direction in the *Unfamiliar* condition (emotional < neutral), our data did not show a significant main effect for *Congruency* (*F*(1, 1001) = .35, *p* = .980) nor an interaction effect between *Congruency* and *Familiarity* (*F*(1, 1001) = .29, *p* = .593, Figure 4, also see Table S9)^29^.

As a control, novel zoo visitors performed the same experiment. We found a main effect of *Congruency* (*F*(1, 9784) = 4.12, *p* = .042) but not for *Stimulus Set (F*(1, 9784) = .01, *p* = .930) nor for the interaction between the two (*F*(1, 9784) = .03, *p* = .865, Table S10)^29^. As such, participants had a faster reaction time to a probe replacing an emotional stimulus (*M* = 434.73, *SE* = 4.05) than to a probe replacing a neutral stimulus (*M* = 437.62, *SE* = 4.05, *t*(9784) = 2.03), regardless of the stimulus set. This is important because it shows that the expressions of the keepers attracted as much attention as the ones from the NimStim set, thus, the null-result in bonobos is unlikely to be attributable to any qualitative characteristics of the stimuli used, at least to the human eye. Finally, a Bayesian analysis confirmed that bonobos have no attentional bias towards human emotional expressions (see Supplements).

### Experiment 3: Humans’ attentional bias towards emotions of conspecifics

Testing whether humans have an attentional bias for emotions of familiar and unfamiliar others, we found a significant main effect of *Congruency* (*F*(1, 16214 = 5.80, *p* < .05) and an interaction effect between *Congruency* and *Familiarity* (*F*(1, 16214 = 5.08, *p* < .05, Figure 4 and Table S13). Simple contrasts showed that participants were significantly faster when a probe replaced an emotional stimulus (*M* = 552.90, *SE* = 3.69) versus a probe replacing a neutral stimulus (*M* = 558.68, *SE* = 3.73) in the *Familiar* condition (*t*(16214) = 3.29, *p* < .01) but not in the *Unfamiliar* condition (*p* = .913)^29^. Importantly, when we performed a similar experiment in a novel group of zoo visitors (see Experiment 2 and Table S10), a significant *Congruency* effect was observed, showing that the null-finding in the unfamiliar condition of Experiment 3 is likely a consequence of the inclusion of emotional expressions of familiar others, that rendered the expressions of unfamiliar people irrelevant^29^.

To follow-up the attentional bias towards emotional expressions of familiar others, we investigated whether emotional intensity further boosted this effect. As expected, the more intense a stimulus was rated, the faster it captured attention (*F*(1, 7178 = 6.55, *p* = .011, Table S14)^29^.

## Discussion

Emotional expressions are pivotal to understanding the internal state of others and predicting their future behavior, and as such, receive privileged access to attention ^11,12,30^. Moreover, emotions often arise in social situations involving close others, yet are rarely studied in this context. The current study investigated the potential link between emotional attention and familiarity with the expressor. It included three main experiments and several control and validation experiments including bonobos and humans; two closely related species. The findings show that in both species, attention is attuned to emotional expressions of conspecifics, especially when these expressions are intense. Crucially, in both species emotional attention is moderated by the familiarity of the expresser, albeit in a different manner; bonobos immediately orient their attention towards emotional scenes depicting unfamiliar others, whereas humans more readily pay attention to emotional expressions of family members or friends. Below we discuss the result of each experiment separately and provide suggestions for further research.

Previous research has shown that bonobos have heightened attention to the emotional expressions of unfamiliar conspecifics^2^, and the current study builds on this research. Specifically, by adding photographs of group mates to the stimulus materials, Experiment 1 showed that bonobos’ attentional bias towards emotional expressions is exclusively present when observing unfamiliar, but not familiar individuals. From a human perspective, this finding may appear counter-intuitive. However, this novel finding is in fact in line with previously conducted behavioral studies in bonobos calling attention to their strong xenophilic tendencies and other-regarding preferences (i.e., bonobos voluntarily help non-group members in obtaining food ^31^; bonobos forego their own food in order to facilitate an interaction with a stranger and prefer them over group members, and help strangers acquire food ^19^. Emotional attention can be driven by the evolutionary relevance of the emotional signal to the observer ^32^. It is thought that for bonobos, socializing with unfamiliar conspecifics is beneficial as it helps them extend their social network. In turn, socializing with unfamiliar others may enhance survival by promoting cooperation among individuals ^31^. Our results support this notion, and extend the literature by showing that the brains of bonobos developed to selectively attend to emotional signals from potentially interesting unfamiliar social partners. We could rule out effects of heightened novelty in the unfamiliar condition, because bonobos on average responded as fast to stimuli of unfamiliar (novel) as of familiar individuals. Thus, the socio-emotional nature of bonobos, combined with the novel individuals in the stimuli, likely drew their attention in this experiment. Still, other moderating factors might have been at play that are worthwhile to discuss.

Our initial prediction was that bonobos would show an attentional bias towards emotional expressions of unfamiliar conspecifics and a similar but dampened bias regarding familiar conspecifics. Nevertheless, we found no such bias and the results even pointed in the other direction, as was also the case when they observed unfamiliar humans. It is possible that when viewing familiar individuals, pre-existing knowledge about those individuals interacts with attentional processes, thereby introducing more variation in what captures attention. Other research indeed suggests that social characteristics of the observer in relation to the observed individual(s) may play a role in how emotions are processed; attention has been shown to be modulated by sex^33^, social bond^34,35^, rank^36^, and kinship ^36^. The current study sample does not allow us to disentangle potential effects of social characteristics. Due to the endangered status of bonobos and their low population in zoos, research access to this species is difficult. Furthermore, separation of individuals is prohibited for ethical reasons and because females dominate in this species, our female subjects did not let the males take part in the study^2^. However, inspection of the two bars representing the familiar and unfamiliar condition in Figure 1A suggests that the inter-individual variance was comparable between these two conditions.

Another possibility for why an attentional bias towards the emotional expressions of familiar conspecifics was not observed may be related to the fact that familiar and unfamiliar conspecifics were shown within the same experiment. The emotional expressions of unfamiliar conspecifics may be of such high relevance for this species, that it rendered biases towards expressions of kith and kin insignificant. We elaborate on this interpretation after discussing Experiment 2, as a similar explanation could be put forward for the results in human participants.

Experiment 2 showed that bonobos did not show an attentional bias towards facial expressions of emotion of humans. When repeating this experiment in human participants, an attentional bias towards emotional expressions was observed. Although the expressions were salient enough for humans to capture attention, they may not have been equally salient for bonobos. Our results fit with earlier findings showing that human participants had an attentional bias towards isolated whole bodily expressions of emotion of chimpanzees and humans, but chimpanzees did not ^37^. However, there is some evidence that apes process emotional expressions of humans similarly as those of conspecifics. A study in orangutans showed that they preferentially looked at the emotional expressions of others as compared to neutral expressions, regardless of whether the expressions were of humans or conspecifics^38^. This suggests that orangutans are sensitive to emotions of another, phylogenetically close species. As these two previous studies tap into a different attentional mechanism than of the one in the current study (i.e. sustained attention versus immediate attention), a future direction for research would be to study how bonobos and humans view each other’s facial expressions in an eye tracking paradigm.

In Experiment 3 using a large heterogeneous community sample, we show that human attention is captured by the emotional expressions of family members and friends, and similar to the bonobo results, the more intense expressions are, the faster they capture attention. Traditionally, emotional attention is studied using stimuli that depict unfamiliar individuals only. For the first time, we show that familiarity with the expressor in terms of their social or familial relationship differentially affects the processing of emotions in humans. Humans have strong affinity with their own social group and often choose to associate with others who are similar to themselves in some respect ^39^. These parochial tendencies are likely to be adaptive, as they bolster cooperation between individuals within the same group ^9^. As such, our results contribute to the existing literature by showing that intergroup bias already presents itself early on in social perception by guiding attention to emotions of socially close others. Interestingly, we did not find evidence for an attention bias towards emotions when participants viewed unfamiliar individuals. Crucially, a control experiment showed that it was the presence of familiar individuals within the same experiment that dampened the focus of attention towards emotional expressions of unfamiliar others. It is possible that the social relevance of the stimuli (i.e. for humans seeing kith and kin) interacts with detecting emotional expressions, prompting stronger activation of attentional and emotional brain mechanisms than when viewing emotions of unfamiliar, less-relevant others. Indeed, according to appraisal theory (e.g. ^40^), the social relevance of stimuli to the observer likely drives attentional mechanisms^41^. This relevance is based on personal goals, values, and needs ^14,42^. In our study, the visitors that participated in the dot-probe task and that were featured in the stimuli were closely bonded individuals and visited the zoo together. It is therefore plausible that these friends or family members were important to the participant performing the task, and thus their emotional expressions captured attention faster than neutral expressions. Combined with an evolutionary perspective describing how humans evolved strong parochial tendencies, our results fit best with appraisal theory (e.g. Lazarus, 2001) of emotion processing. To study the effects of social relevance on the recruitment of attentional mechanisms, an interesting future direction would be to investigate how the degree of social closeness (e.g. familiar friend versus a familiar non-friend) affects emotional attention.

To conclude, our study contributes to the understanding of how evolution shaped other-regarding preferences of bonobos and humans by showing that they are deeply ingrained in early social perception and, crucially, are shared between the species. At the same time the results demonstrate that bonobos and humans show clear differences in attention for emotional signals of familiar compared to unfamiliar others, despite the shared underlying attentional mechanisms. Our findings support appraisal theories of emotion that suggest that the personal relevance of signals to observers moderates attention for emotions. Moreover, our results support an evolutionary perspective on emotion processing that suggests that the last common ancestor of humans and bonobos likely shared a similar attentional mechanism for guiding attention towards emotions. Nevertheless, differences in the environments of bonobos and humans likely helped shape the striking differences in how bonobos and humans perceive emotions of familiar and unfamiliar others. Moreover, as a recent review points out, emotional expressions are multifaceted and while there are key similarities between human and great ape expressions of emotions, there are also species-specific elements that require more thorough comparison ^24^. For instance the highly expressive eyebrows in humans, the white sclera surrounding the iris and the contrast between skin color and redder lips ^24^. To understand both homologies and divergences within emotional expressions in humans, bonobos and other great apes, future studies could benefit from complementary tools such as eye tracking to specifically investigate sustained attention for emotional expressions of group members or strangers.

## Acknowledgements

We thank the Kyoto University Primate Research Institute for donating an automatic feeder and Coos Hakvoort for constructing the bonobo setup. We thank Keetie de Koeier, Iris Schapelhouman, Grietje Grootenhuis, Jacqueline Ruijs, and Carolijn de Jong for their crucial assistance during testing. Additionally, we thank Daan Laméris for coding the videos and Berta Torres for scoring stimuli. Finally, we thank all students who helped in collecting the human data.

