## Supplemental Materials for "Attention Towards Emotions is Modulated by Familiarity with the Expressor. A Comparison Between Bonobos and Humans"

**EXPERIMENT 1** ........................................ 3-8
**Supplemental methods**........................................ 3-7
*Statistical analysis for control experiment*........................................ 12-13
**Supplemental results**........................................ 14-15
*Bayesian analysis*........................................ 15

**EXPERIMENT 3**........................................ 16-20
**Supplemental methods**........................................ 16-19
*Participants*........................................ 16-17
*Stimulus validation*........................................ 17-18

**Experiment 1: Bonobos' attentional bias towards emotions of conspecifics.**

### SUPPLEMENTAL METHODS

**Participants.** The following individual bonobos were tested: Besede, 11 years old; Monyama, 6 years old; Kumbuka, 17 years old; Yahimba, 7 years old and daughter of Kumbuka. At time of testing, the group consisted of 12 individuals (3 males) housed in large in- and outdoor enclosures (2970 $m^2$ in total) containing several climbing structures, trees, bushes and ropes, puzzles from which they could acquire food, and small streams of water. To mimic natural fission-fusion behavior, bonobos were always housed in two separated groups that varied in composition regularly. Non-participating bonobos were distracted by the caretaker with a body-part training in which bonobos were instructed to present specific body parts to the animal caretaker, and were rewarded with an apple cube for each correct presentation, just like the participating bonobos when they completed a trial. All participants in this study were exposed to humans since birth and interacted with them on a daily basis. Daily diet consisted of a variety of fruits, vegetables, branches and leaves, and pellets enriched with necessary nutrients. The bonobos were fed four to five times a day, and water was available ad libitum. Furthermore, bonobos were never deprived of water or food at any stage of the experiment.

**Stimulus material.** While we currently do not fully understand what bonobo emotions exactly entail, we rely on existing observational work to define emotional states. Using Anderson and Adolphs' (2014) componential view of emotions in animals, we use socio-emotional scenes of bonobos engaged in sex, play, grooming, and bonobos scratching, yawning (indicators of stress [1]) or in distress (e.g. silent bared-teeth display) as reflections of emotional states. We used similar emotion categories as Kret et al. (2016)[2] (but all novel images), with the exception that we included scratching as a new category and left out pant hoot and food, because they did not attract attention over neutral scenes in our previous study.

**Table S1.** Number of pictures per familiarity category and per emotion category for bonobos in Experiment 1.

| | Familiarity | |
|---|---|---|
| Picture category | Unfamiliar | Familiar |
| Distress | 15 | 25 |
| Sex | 27 | 16 |
| Play | 22 | 36 |
| Groom | 53 | 50 |
| Yawn | 20 | 16 |
| Scratch | 18 | 30 |
| Total emotional | 155 | 173 |
| Total neutral | 155 | 173 |

*Note.* All bonobo participants saw the same number of unique pictures, but for the familiar models the composition of the stimulus set differed per participant because we replaced pictures of the participants themselves with pictures of other familiar individuals. Furthermore, the number of pictures differs per *Familiarity* and also per *Picture category* because some behaviours were easier to photograph or occurred more frequently than other behaviours.

**Table S2.** Definition of emotion categories used for bonobos in Experiment 1.

| Picture category | Description | No. of individuals per picture |
|---|---|---|
| Distress | Aggressive displays, such as long and short charges, mutual and direct displays; submissive behaviours such as grin faces, fleeing, and crouching | ($M$ = 1.55, $SE$ = 0.16) |
| Sex | mating, genito-genital rubbing, masturbation, prominent full swelling, penile erection | ($M$ = 2.25, $SE$ = 0.25) |
| Playing | Together or alone, with a relaxed open mouth, without an object | ($M$ = 2.25, $SE$ = 0.25) |
| Grooming | Two or more individuals grooming | ($M$ = 2.38, $SE$ = 0.26) |
| Yawning | Wide open mouth with or without canine visibility | ($M$ = 1.17, $SE$ = 0.17) |
| Scratching | Fingers separated and touching the body, face, or one of the limbs | ($M$ = 1.00, $SE$ = 0.00) |
| Neutral | Walking, lying down, or sitting | ($M$ = 1.67, $SE$ = 0.14) |

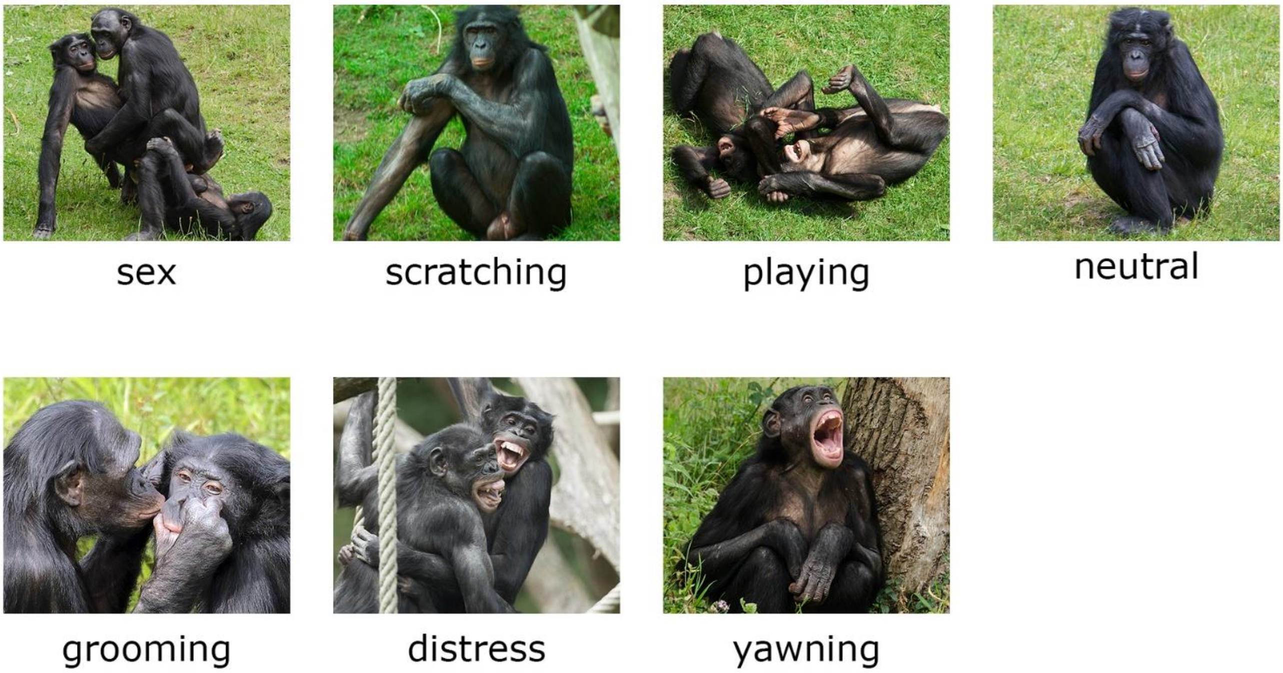

**Figure S1.** Examples of stimuli of all emotional categories used in Experiment 1. An emotional picture was always paired with a neutral picture. The emotional and neutral pictures were of either familiar or unfamiliar individuals.

**Stimulus validation.** Six primate experts scored the pictures of bonobos based on valence and intensity. Three experts worked with the bonobos on a daily basis, and the others worked with bonobos or chimpanzees in the past. Experts were presented with one picture at a time and were asked to 1) determine how negative or positive they thought bonobos would experience each picture (1= very negative, 7 = very positive) and 2) how intense the picture was (1= not intense, 7 = very intense). Pictures were shown until a response was given. We calculated intraclass correlations (ICCs) for valence and intensity ratings using a two-way mixed model and a consistency definition and found a high reliability for both ratings ($ICC_{valence}$ = .82, 95% CI: .79 - .84, $F(653, 3265) = 5.45$, $p < .001$; $ICC_{intensity}$ = .87, 95% CI: .86-.89, $F(653, 3265) = 7.96$, $p < .001$). A generalized linear mixed model with *Emotionality* (emotion versus neutral) as fixed factor, *Rating* as target variable and *Rater Number* as random effect confirmed that emotional pictures are indeed rated higher in intensity than neutral pictures ($M_{emotional} = 3.28$, $SE = 0.35$, $M_{neutral} = 1.76$, $SE = 0.18$, $F(1, 3922) = 1337.81$, $p < .001$).

**Table S3.** Overview of average intensity and valence ratings per emotion category and per familiarity category in Experiment 1.

| | Familiar | | Unfamiliar | |
|---|---|---|---|---|
| Emotion category | Average Valence (SD) | Average Intensity (SD) | Average Valence (SD) | Average Intensity (SD) |
| Distress | 3.07 (1.38) | 4.73 (1.57) | 3.49 (1.55) | 4.74 (1.39) |
| Sex | 5.93 (0.85) | 4.90 (1.28) | 5.88 (0.93) | 4.94 (1.34) |
| Play | 5.31 (0.95) | 3.12 (1.58) | 5.44 (0.92) | 2.89 (1.44) |
| Groom | 5.31 (0.82) | 3.09 (1.64) | 5.15 (0.83) | 2.73 (1.57) |
| Yawn | 3.96 (0.92) | 3.31 (1.56) | 3.97 (1.00) | 3.53 (1.62) |
| Scratch | 4.11 (1.07) | 1.76 (0.71) | 4.14 (1.02) | 1.74 (0.78) |
| Neutral | 4.63 (0.91) | 1.64 (0.89) | 4.62 (0.88) | 1.74 (0.98) |

**Procedure.** The bonobos were already familiarized with the dot-probe procedure during a previous study [2]. To make sure they were still capable of conducting the task, we started with a two-month training period (about 7 sessions per ape) in which they performed a dot-probe task with pictures of black rabbits and goats. Only after all the apes were able to correctly pass 95% or more of the trials within one session, we moved on to the experiment. To start a trailing or experimental session, we called forth the highest-ranking participating individual of the subgroup that was present in the enclosure. The other bonobos were distracted by the caretaker.

**Table S4.** Number of incorrect trials per bonobo on the dot-probe with bonobo stimuli in Experiment 1.

| Participant name | Tested trials (sessions) | Repetitions of trials* | Incorrect trials (% of grand total) † | Of which due to nose wipes | Of which due to scratching | Of which due to outliers‡ |
|---|---|---|---|---|---|---|
| Besede | 525 (21) | 202 | 130 (6.0 %) | 2 | 1 | 127 |
| Kumbuka | 582 (24) | 264 | 181 (8.4%) | 38 | 2 | 141 |
| Monyama | 537 (22) | 212 | 104 (4.8%) | 2 | 0 | 102 |
| Yahimba | 518 (21) | 210 | 99 (4.6%) | 4 | 0 | 95 |
| Grand total | 2162 | 888 | 514 (23.8 %) | 46 | 3 | 465 |

* These reflect the number of trials that were repeated due to disruptions in the first/original trials.

† The number of incorrect trials consists of both erroneous trials within the first/original trials and erroneous trials within the repetitions.

‡ Outliers were disruptions during a trial other than due to the behaviours described above, e.g.: not attending to the screen during stimulus presentation, someone other than the participant pressing the probe, using the opposing hand, not sitting directly in front of the screen, or the screen not immediately responding to a touch. Outliers also contain extreme RTs (250 < RT < 5000) and RTs higher than the median RT per participant minus 2.5*MAD per participant.

**Data filtering.** Bonobos were not always paying attention to the screen during the task. We considered several behaviors as reasons to discard a trial from further analyses: e.g. bonobos did not immediately touch the probe, were distracted by other bonobos or did not attend to the screen, when another individual pressed the probe, or when hands were switched within a trial, or when bonobos performed movements that interfered with the task (scratching or nose wiping).

**SUPPLEMENTAL RESULTS**

**Table S5.** Bonobo results on an attentional bias towards emotional expressions versus neutral expressions of familiar and unfamiliar bonobos in Experiment 1.

| Source | *F* | df1 | | df2 | | *P* value |
|---|---|---|---|---|---|---|
| Congruency (*emotional*, *neutral*) | 4.65 | 1 | | 164 | | .031 |
| Familiarity (*familiar, unfamiliar*) | 0.00 | 1 | | 164 | | .959 |
| Congruency*Familiarity | 4.654 | 1 | | 164 | | .031 |
| Random effects | Estimate | SE | *Z* | *P value* | 95% confidence interval | |
| Variance | 0.02 | 0.00 | 28.61 | .000 | 0.019 | 0.022 |
| Intercept (ID) | 0.05 | 0.04 | 1.22 | .224 | 0.010 | 0.254 |
| Intercept (ID*Session) | 0.00 | 0.00 | 1.17 | .243 | 0.000 | 0.003 |

**Table S6.** Bonobo results on the effect of emotional intensity on attentional bias in stimuli depicting unfamiliar bonobos in Experiment 1.

| Source | *F* | df1 | | df2 | | *P* value |
|---|---|---|---|---|---|---|
| *Intensity Difference Score** (scale) | 8.38 | 1 | | 859 | | .004 |
| Random effects | Estimate | SE | *Z* | *P value* | 95% confidence interval | |
| Variance | 0.02 | 0.00 | 20.63 | .000 | 0.019 | 0.023 |
| Intercept (ID) | 0.05 | 0.04 | 1.22 | .224 | 0.010 | 0.250 |
| Variance (ID * Session) | 0.00 | 0.00 | 0.90 | .368 | 0.000 | 0.004 |

* This difference score refers to the difference in emotional intensity of the probe image compared to the distractor (nonprobe) image.

**Experiment 2: Bonobos' attentional bias towards emotions of humans.**

**SUPPLEMENTARY METHODS**

**Stimulus material.** Emotional and neutral pictures were taken of the face of all 6 familiar individuals with a neutral, white background, and were contrasted with emotional and neutral pictures of 6 unfamiliar, Caucasian female models from the NimStim Set of Facial Expressions [3]. While making the photos of the caretaker, the experimenter enacted the facial expressions as an example for the caretakers and instructed them to mimic her. Photos of emotional expressions were taken in the following order: anger, fear, happiness, sadness, surprise, disgust. Each photo of an emotional expression was followed by a photo of a neutral expression to ensure that neutral photos were slightly different from each other. If the experimenter thought a photo was not similar enough to the faces used from the NimStim database, the photo was retaken. Stimuli from familiar individuals were all unique. However, because the NimStim database sometimes contained only two different neutral expressions per model, for the unfamiliar stimuli only the emotional pictures were all unique.

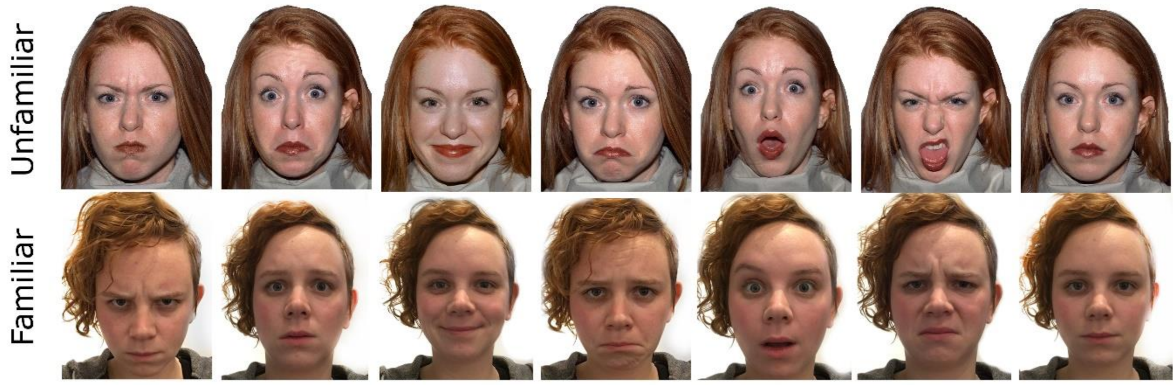

**Figure S2.** Examples of the stimuli of emotional and neutral expressions of unfamiliar (NimStim Faces, up) and familiar (i.e. caretaker or experimenter, below) individuals. The NimStim models were females 1, 2, 3, and 7, 8, 9 [3]. Emotional categories are (left to right): anger, fear, happiness, sadness, surprise, disgust, and neutral. An emotional picture was always paired with a neutral picture, and these pictures were always either from the NimStim or caretaker stimulus set.

**Stimulus validation.**

**Table S7a**. Overview of average agreement on emotionality (emotional/neutral), average rating scores for emotional intensity and authenticity rated by N=5 research assistants in Experiment 2.

| | Caretaker ("familiar") | | | NimStim ("unfamiliar") | | |
|---|---|---|---|---|---|---|
| Stimulus category | Agreement on emotionality (SD) | Intensity rating (SD) | Authenticity rating (SD) | Agreement on emotionality (SD) | Intensity rating (SD) | Authenticity rating (SD) |
| Emotional | 92.8% (.26) | 4.21 (1.36) | 4.37 (1.60) | 88.9% (.32) | 5.09 (1.34) | 3.66 (1.56) |
| Neutral | 77.3% (.42) | 2.84 (1.67) | 5.04 (1.32) | 100% (.00) | 3.18 (1.58) | 4.80 (1.39) |

*Note.* Agreement on emotion refers to the average agreement between 5 raters on whether emotional stimuli were recognized as an emotion by them, and neutral stimuli as neutral. Agreement on whether a stimulus was emotional or neutral was not 100% on average, meaning raters sometimes rated a neutral expression as emotional and vice versa. Furthermore, low intensity scores for neutral stimuli and high intensity scores for emotional stimuli are preferred, because this means there is a clear discrepancy between two simultaneously presented stimuli and as such, highly intense (emotional) stimuli can capture attention faster than low-intensity (neutral) stimuli. A generalized linear mixed model with *Emotionality* (emotion versus neutral) as fixed factor, *Rating* as target variable and *Rater Number* as random effect confirmed that emotional pictures are indeed rated higher in intensity than neutral pictures ($M_{\text{emotional}} = 4.70$, $SE = 0.39$, $M_{\text{neutral}} = 2.79$, $SE = 0.24$, $F(1, 603) = 233.19$, $p < .001$).

**Table S7b.** ICCs of scores on the different scales (intensity, emotionality (emotion/neutral), and authenticity using two-way mixed effects using a consistency definition in Experiment 2 (N=6).

| Scale | Intraclass Correlation | 95% confidence interval | | *F* value (df1, df2) | *P* value |
|---|---|---|---|---|---|
| Intensity | .78 | .71 | .84 | 4.53 (120, 480) | .000 |
| Emotion | .66 | .55 | .75 | 2.94 (120, 480) | .000 |
| Authenticity | .69 | .59 | .77 | 3.20 (120, 480) | .000 |

**Procedure.**

**Table S8.** Number of incorrect trials per bonobo on the dot-probe with human stimuli Experiment 2.

| Participant name | Tested trials (sessions) | Repetitions of trials* | Incorrect trials (% of grand total) † | Of which due to nose wipes | Of which due to scratching | Of which due to outliers‡ |
|---|---|---|---|---|---|---|
| Besede | 325 (13) | 198 | 66 (20.3) | 0 | 0 | 58 |
| Kumbuka | 326 (13) | 198 | 106 (32.5) | 15 | 0 | 73 |
| Monyama | 377 (15) | 245 | 78 (20.7) | 0 | 0 | 48 |
| Yahimba | 350 (14) | 223 | 123 (35.1) | 1 | 0 | 58 |
| Grand total | 1378 | 864 | 373 | 16 | 0 | 237 |

* These reflect the number of trials that were repeated due to disruptions in the first/original trials.

† The number of incorrect trials consists of both erroneous trials within the first/original trials and erroneous trials within the repetitions.

‡ Outliers were disruptions during a trial other than due to the behaviours described above, e.g.: not attending the screen during stimulus presentation, someone other than the participant pressing the probe, using the opposing hand, not sitting directly in front of the screen, or the screen not immediately responding to a touch. Outliers also contain extreme RTs (250 < RT < 5000) and RTs higher than the median RT per participant minus 2.5*MAD per participant.

**Methods for human participants performing the control experiment**

**Stimuli for control experiment.** We created two versions of the task each containing 72 trials with 36 trials of "familiar" humans and 36 "unfamiliar" humans. The only difference between the two versions was that the probe location was mirrored (i.e. if in version 1 it appeared behind one of the emotional pictures, it appeared behind the neutral stimulus in version 2, and vice versa). None of the human participants knew the "familiar" humans on the stimuli. This was intentional; the aim was to control for potential differences in the two stimulus sets (i.e. NimStim ("unfamiliar") on the one hand and caretakers ("familiar") on the other) and help explain the bonobo results. Per participant, every stimulus was only shown once.

**Apparatus for control experiment.** We set up a table near the bonobo enclosure (Figure 2 in main text). Here, participants performed the experiment on an Iiyama ProLite T1930SR1 touchscreen (19", 1280x1024 pixels, ISO 5ms), using E-Prime 2.0. The touchscreen was placed on a table and participants were seated with their back against a wall to prevent others from distracting them.

**Procedure for control experiment.** Human participants were recruited by actively approaching them when they passed by the bonobo enclosure. They were told the bonobos participated in a computerized task and were asked whether they were interested in conducting the same task. If they agreed to participate, they were given an information sheet explaining the procedure of the task and a consent form to sign. The participant then took place at the table behind a touchscreen and the experimenter started the dot probe task. Because bonobos could not receive instructions, we also decided to keep instructions for the task for human participants to a minimum. The task starts with pictures of the four participating bonobos with the message "Are you faster than the bonobos? Tap the dot on the screen as soon as it appears. To continue to a brief training session, click anywhere on the screen". Next, participants went through 3 practice trials with flowers as stimuli, and subsequently started with the experiment. The experiment was exactly the same as the bonobo dot-probe task from Experiment 2, except that humans completed 72 trials. After finishing the task, participants could see their average reaction time and how it compared to the average reaction time of the bonobos. They were then debriefed about the study and thanked for their participation.

**Data filtering for control experiment.** For human participants, we first divided every participant into an age category as reaction times can be higher in older versus younger individuals (i.e. 18-20, 21-25, 26-30, etc.). Next, we filtered out extreme RTs (>5000 ms) and then calculated the median absolute deviation for reaction times per age category. Finally, we used the following data filter: [RT < (Median RT + (2.5 * Mean Absolute Deviation))]. 606 Trials (5.61%) were subsequently removed for further analysis.

**Statistical analysis for control experiment.** The human data assist in and strengthen interpretations of the bonobo data, and therefore we had the following hypotheses: I) humans have an attentional bias towards emotional expressions of other humans versus neutral expressions, and II) there are no differences in attention for the emotional or neutral expressions between the two stimulus sets (NimStim stimulus set ("unfamiliar") versus caretaker stimulus set ("familiar")). In the analysis, we included *Congruency* and *Stimulus set* (caretaker stimuli and NimStim stimuli) and their interaction as fixed factors, and reaction time as the target variable. We used a nested structure with participants (150) nested within trials (72), and for the target variable we used a gamma-distribution with a log-link function.

**SUPPLEMENTAL RESULTS**

**Table S9.** Bonobo results on an attentional bias towards emotional expressions versus neutral expressions of familiar (caretaker) and unfamiliar (NimStim) humans in Experiment 2.

| Source | *F* | df1 | | df2 | | *P value* |
|---|---|---|---|---|---|---|
| Congruency (*emotional*, *neutral*) | .35 | 1 | | 1001 | | .553 |
| Familiarity (*familiar, unfamiliar*) | .00 | 1 | | 1001 | | .980 |
| Congruency*Familiarity | .29 | 1 | | 1001 | | .593 |
| Random effects | Estimate | SE | *Z* | *P value* | 95% confidence interval | |
| Variance | 0.03 | 0.00 | 22.25 | .000 | 0.024 | 0.029 |
| Intercept (ID) | 0.04 | 0.03 | 1.16 | .246 | 0.006 | 0.189 |
| Variance (ID * Session) | 0.01 | 0.00 | 1.83 | .067 | 0.002 | 0.015 |

**Table S10.** Human results on an attentional bias towards emotional expressions versus neutral expressions of other human stimuli involving the caretaker versus NimStim stimulus set Experiment 2.

| Source | *F* | df1 | | df2 | | *P value* |
|---|---|---|---|---|---|---|
| Congruency (*emotional*, *neutral*) | 4.12 | 1 | | 9784 | | .042 |
| Stimulus set (C*aretaker, NimStim)* | .01 | 1 | | 9784 | | .930 |
| Congruency*Stimulus set | .03 | 1 | | 9784 | | .865 |
| Random effects | Estimate | SE | *Z* | *P value* | 95% confidence interval | |
| Variance | 0.03 | 0.00 | 69.40 | .000 | 0.025 | 0.027 |
| Intercept (ID) | 0.01 | 0.00 | 8.31 | .000 | 0.010 | 0.015 |

**Bayesian analysis to confirm that bonobos do not have an attentional bias for human emotional expressions in Experiment 2.**

We performed a Bayesian Generalized Linear Model analysis using the brms package [4] in R. In the model, *Congruency*, *Familiarity* and their interaction were defined as fixed factors and *Participant*Session* as random factor, and we used a gamma distribution and log-link function with the standard priors of brms and based on a tutorial [5]. We compared this model to the null-model (with only the random factor) and found a logarithmic Bayes factor of $logBF_{01} = -19.79$, indicating strong evidence for the null-hypothesis over the alternative hypothesis [6]. Thus, given our data, we found no evidence that bonobos have an attentional bias towards human facial expressions of emotion.

**Experiment 3: Humans' attentional bias towards emotions of conspecifics.**

**Participants.**

**Table S11a.** Descriptives of participants and their relation to the participant on the stimuli in Experiment 3.

| Relationship participant on task versus participant on photo* | Sex participant on task versus Sex participant on photos | | | | |
|---|---|---|---|---|---|
| | Male versus Male | Male versus Female | Female versus Female | Female versus Male | Grand total |
| Brother/sister | 25 | 25 | 32 | 19 | 101 |
| Child | 25 | 16 | 33 | 23 | 97 |
| Parent | 22 | 23 | 25 | 28 | 98 |
| Spouse/partner | 1 | 27 | 0 | 48 | 76 |
| Niece/nephew† | 0 | 1 | 1 | 0 | 2 |
| Friend/Colleague | 8 | 3 | 12 | 7 | 30 |
| Grand total | 81 | 95 | 103 | 125 | 404‡ |

* The relationship is seen from the viewpoint of the participant doing the task, e.g. "Child" means that the stimuli are of the child of the participant who performed the dot probe task.

† As we were interested in how closely bonded individuals attend to each other's emotions, we focused mainly on families and friends. We did not collect a lot of participants with a more distant family relationship (e.g. aunts/uncles, nephews/nieces, cousins), but decided not to remove the 2 participants with a niece or nephew.

‡ Note that this number is not the same as the one reported in the main text (N=449), this is because of a technical failure, the relationship data of 45 participants was not registered.

**Table S11b.** Information on how often participants in the task saw each other in Experiment 3.

| Relationship participant on task versus participant on photo* | Contact frequency between participant on task and photo | | | |
|---|---|---|---|---|
| | Daily | Weekly | Monthly | Yearly |
| Brother/sister | 88 | 6 | 5 | 2 |
| Child | 92 | 3 | 2 | 0 |
| Parent | 87 | 6 | 5 | 0 |
| Spouse/partner | 50 | 25 | 1 | 0 |
| Niece/nephew† | 1 | 0 | 1 | 0 |
| Friend/Colleague | 13 | 9 | 7 | 1 |
| Grand total | 331 | 49 | 21 | 3 |

**Stimulus validation.**

**Table S12a**. Overview of average agreement on emotionality (emotion/neutral), and average rating scores for emotional intensity and authenticity of human stimuli in Experiment 3.

| Rater group | Neutral stimuli | | | Emotional stimuli | | |
|---|---|---|---|---|---|---|
| | Agreement on emotionality (SD) | Intensity rating (SD) | Authenticity rating (SD) | Agreement on emotionality (SD) | Intensity rating (SD) | Authenticity rating (SD) |
| 1 | 82.9% (.38) | 3.16 (1.85) | 5.72 (2.41) | 68.3% (.47) | 4.08 (1.86) | 4.85 (1.87) |
| 2 | 74.8% (.43) | 2.17 (1.64) | 5.79 (1.42) | 88.3% (.32) | 4.97 (1.79) | 5.39 (1.65) |
| 3 | 77.3% (.42) | 2.69 (1.3) | 5.26 (1.66) | 90.5% (.29) | 4.44 (1.59) | 4.06 (1.88) |
| Grand average | 81.5% (.39) | 2.99 (1.83) | 5.71 (1.29) | 70.9% (.45) | 4.18 (1.71) | 5.29 (1.43) |

*Note.* Rater group 1, N = 8; Group 2, N = 5; Group 3, N = 5. Agreement on emotion/neutral refers to the average agreement between raters on whether emotional stimuli were recognized as an emotion by them, and neutral stimuli as neutral. Agreement was not 100% on average, meaning raters sometimes rated an emotional expression as neutral and vice versa. Intensity and authenticity were rated on a scale from 1-7 (1 = low intensity/authenticity, 7 = high intensity/authenticity). Furthermore, low intensity scores for neutral stimuli and high intensity scores for emotional stimuli are preferred, because this means there is a clear discrepancy between two simultaneously presented stimuli and as such, highly intense (emotional) stimuli can capture attention faster than low-intensity (neutral) stimuli. A generalized linear mixed model with *Emotionality* (emotion versus neutral) as fixed factor, *Rating* as target variable and *Rater Group*Rater Number* as random effect confirmed that emotional pictures are indeed rated higher in intensity than neutral pictures ($M_{emotional}$ = 4.17, $SE$ = 0.22, $M_{neutral}$ = 2.74, $SE$ = 0.14, $F(1, 30377) = 5139.47$, $p < .001$).

**Table S12b.** ICC (two-way mixed, consistency) on intensity scores per rater group in Experiment 3.

| Rater group | Intraclass Correlation | 95% confidence interval | | F value (df1, df2) | *P* value |
|---|---|---|---|---|---|
| 1 | .74 | .73 | .76 | 3.90 (3078, 21546) | .000 |
| 2 | .89 | .88 | .91 | 9.34 (568, 2748) | .000 |
| 3 | .78 | .74 | .82 | 4.63 (263, 1052) | .000 |

**Table S12c.** ICC (two-way mixed, consistency) on ratings of emotion type per rater group in Experiment 3.

| Rater group | Intraclass Correlation | 95% confidence interval | | F value (df1, df2) | *P* value |
|---|---|---|---|---|---|
| 1 | .92 | .91 | .92 | 12.07 (3219, 22533) | .000 |
| 2 | .69 | .65 | .72 | 3.18 (687, 2748) | .000 |
| 3 | .82 | .78 | .85 | 5.49 (263, 1052) | .000 |

**Table S12d.** ICC (two-way mixed, consistency) on authenticity scores per rater group in Experiment 3.

| Rater group | Intraclass Correlation | 95% confidence interval | | F value (df1, df2) | *P* value |
|---|---|---|---|---|---|
| 1 | .70 | .68 | .71 | 3.29 (3078, 21546) | .000 |
| 2 | .60 | .55 | .64 | 2.47 (687, 2748) | .000 |
| 3 | .74 | .69 | .79 | 3.87 (263, 1052) | .000 |

**Procedure.** When creating the stimulus material for the dot probe with zoo visitors, the experimenter asked the photo participant to express one of the four emotions (anger, fear, happiness, sadness) based on an example from Model 1 from the NimStim database printed on a sheet of paper. Between each photograph of an emotional expression, the experimenter also took a photo of a neutral expression. Low-quality photos were retaken on the spot.

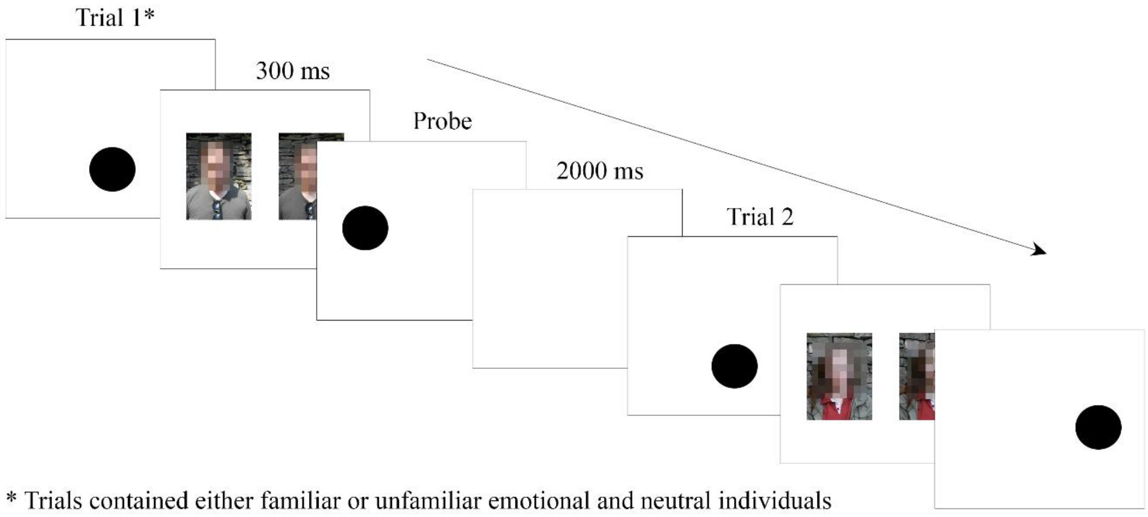

**A**

**Figure S3.** Trial outline for the human dot-probe in Experiment 3 (and the control experiment in Experiment 2). Similar to the bonobo study, the task started with a start-dot, followed by the presentation of an emotional and neutral stimulus of a familiar or unfamiliar individual, of which one was replaced with the probe. The inter-trial interval was 2000ms.

**SUPPLEMENTARY RESULTS**

**Table S13.** Human results on an attentional bias towards emotional expressions versus neutral expressions of familiar and unfamiliar humans in Experiment 3.

| Source | *F* | df1 | | df2 | | *P value* |
|---|---|---|---|---|---|---|
| Congruency (*emotional*, *neutral*) | 5.80 | 1 | | 16214 | | .016 |
| Familiarity (*familiar, unfamiliar*) | 0.04 | 1 | | 16214 | | .834 |
| Congruency*Familiarity | 5.08 | 1 | | 16214 | | .024 |
| Random effects | Estimate | SE | *Z* | *P value* | 95% confidence interval | |
| Variance | 0.02 | 0.00 | 88.76 | .000 | 0.020 | 0.021 |
| Intercept (ID) | 0.02 | 0.00 | 14.30 | .000 | 0.015 | 0.020 |

**Table S14.** Results of testing whether a difference score in intensity between the probe image and nonprobe image affected reaction times in humans in Experiment 3.

| Source | *F* | df1 | | df2 | | *P* value |
|---|---|---|---|---|---|---|
| *Intensity Difference Score* (scale) | 6.55 | 1 | | 7178 | | .011 |
| Random effects | Estimate | SE | *Z* | *P value* | 95% confidence interval | |
| Variance | 0.02 | 0.00 | 58.17 | .000 | 0.020 | 0.021 |
| Intercept (ID) | 0.018 | 0.00 | 12.97 | .000 | 0.015 | 0.021 |
